## Supplementary figures and images for "Comparative whole-genome analyses of articular chondrocytes and skin fibroblasts reveal distinct genome instability landscapes in mesenchymal cell types"

### Fig S1.tif

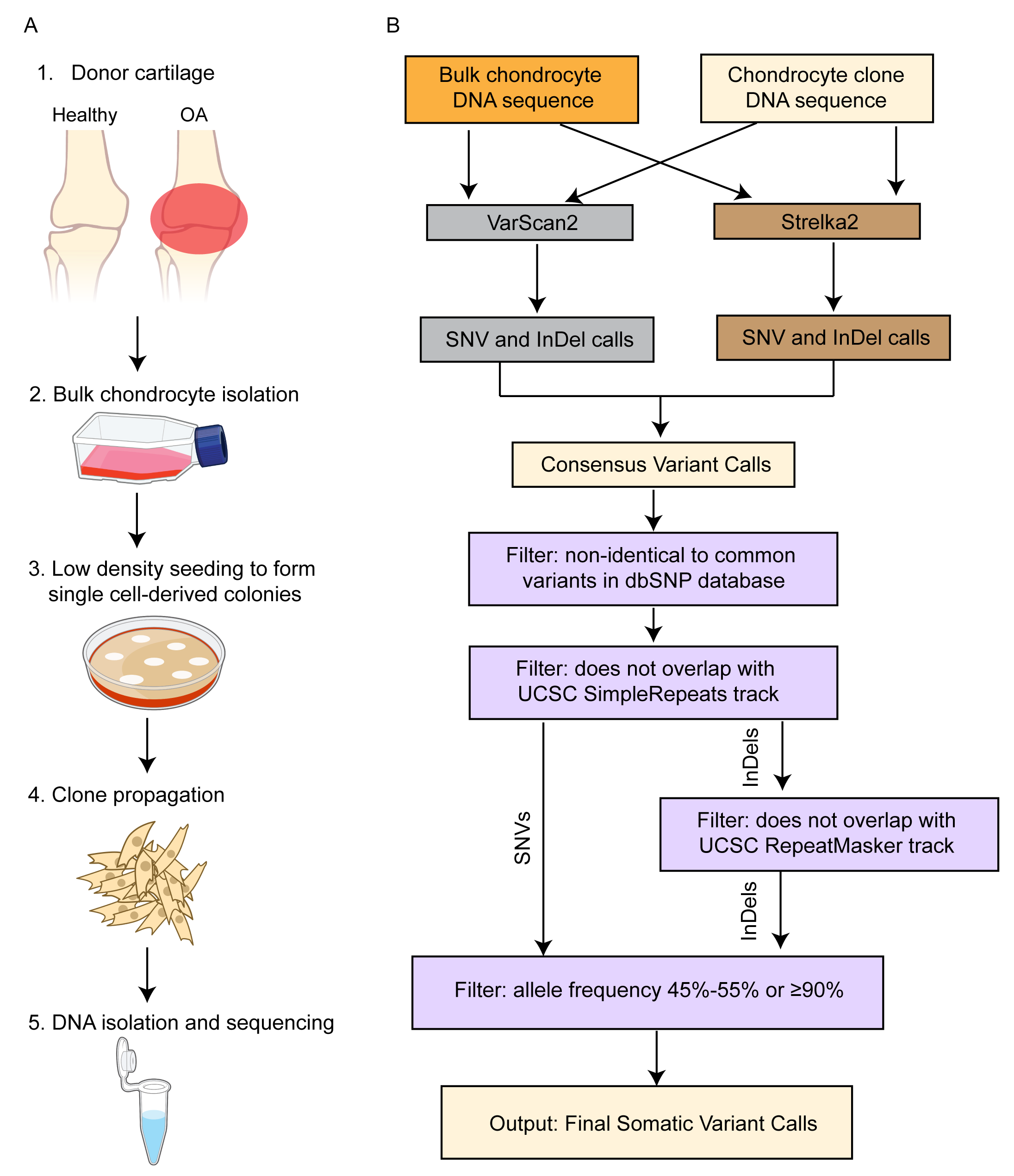

### Fig S2.tif

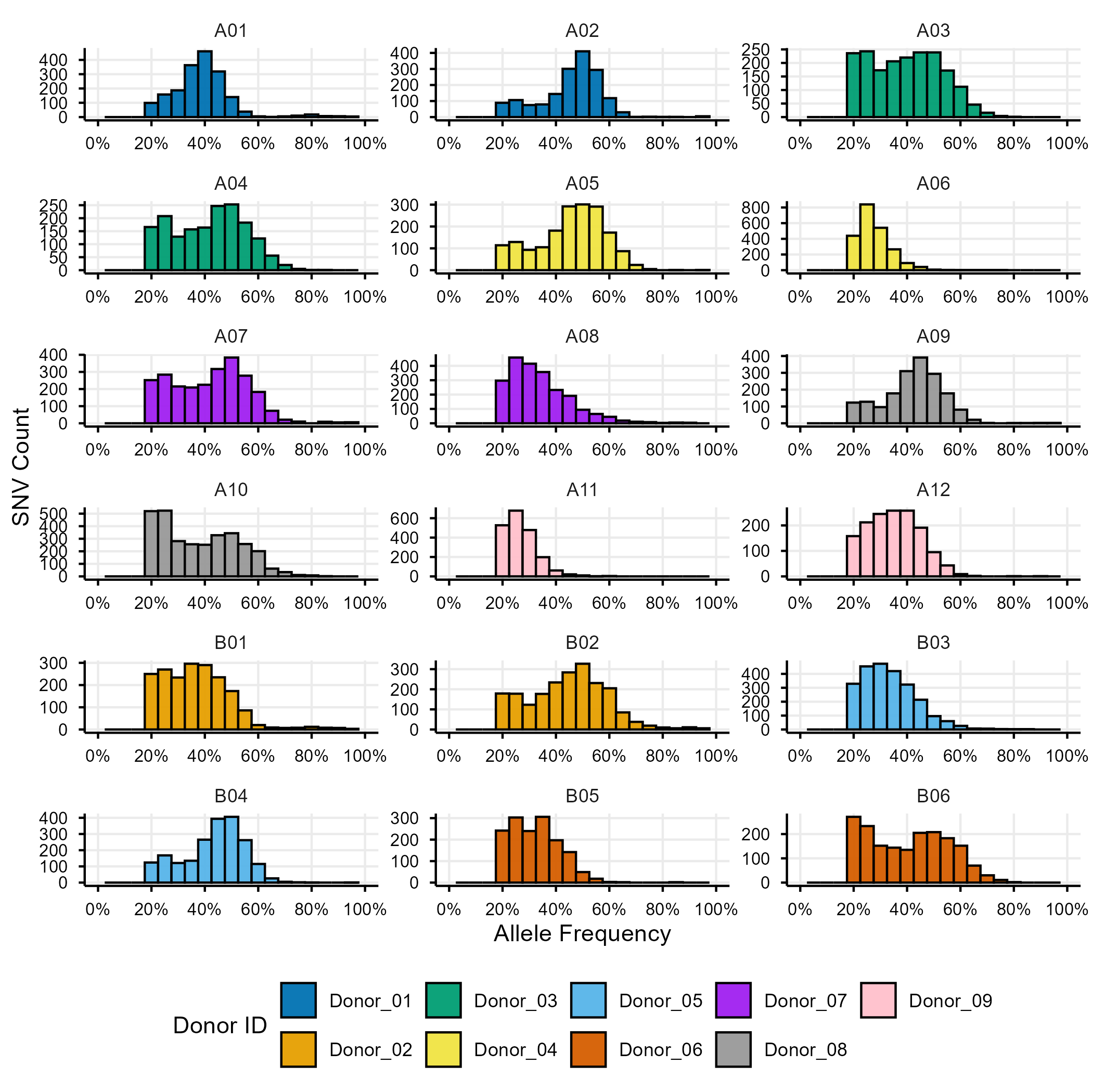

### Fig S3.tif

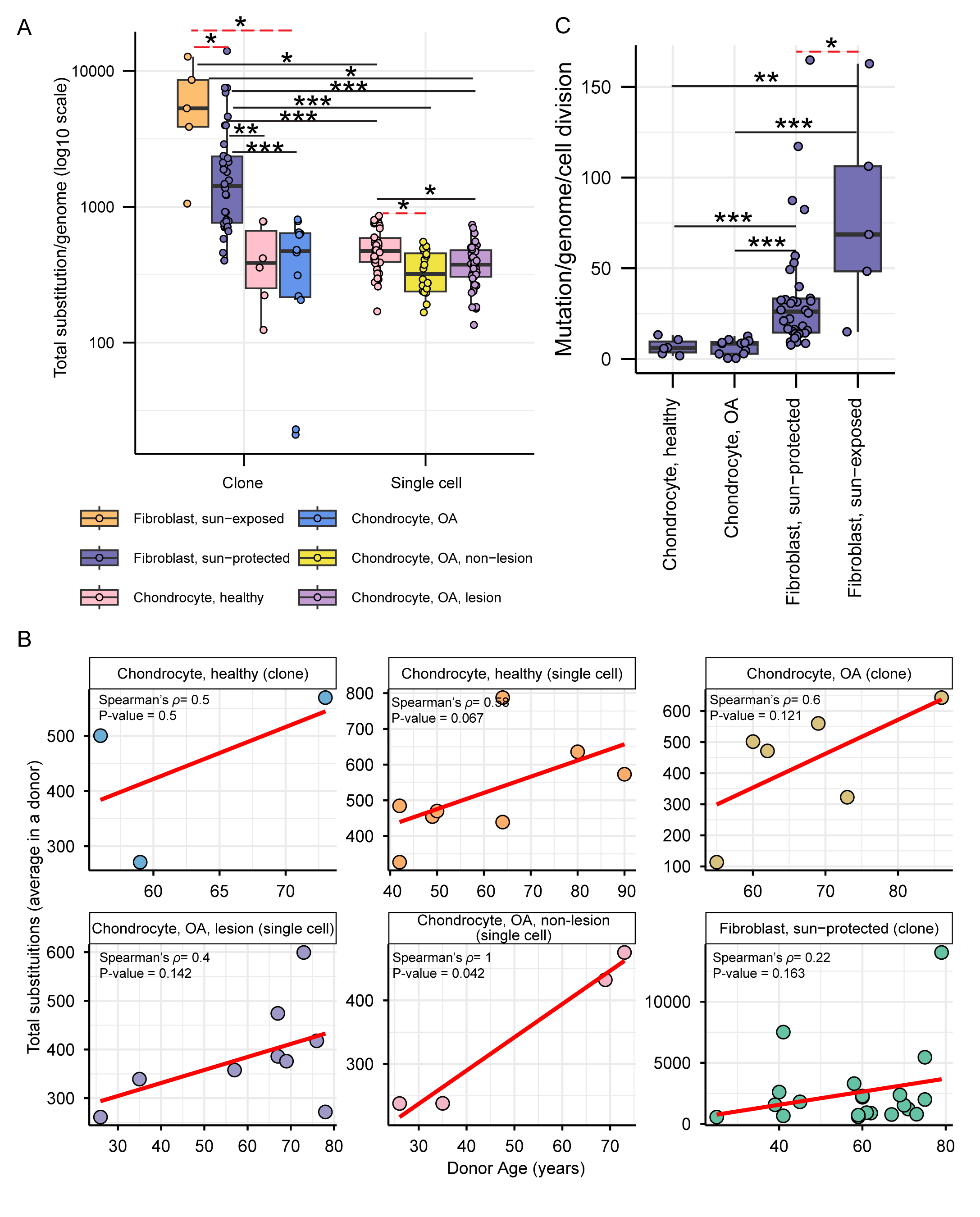

### Fig S4.tif

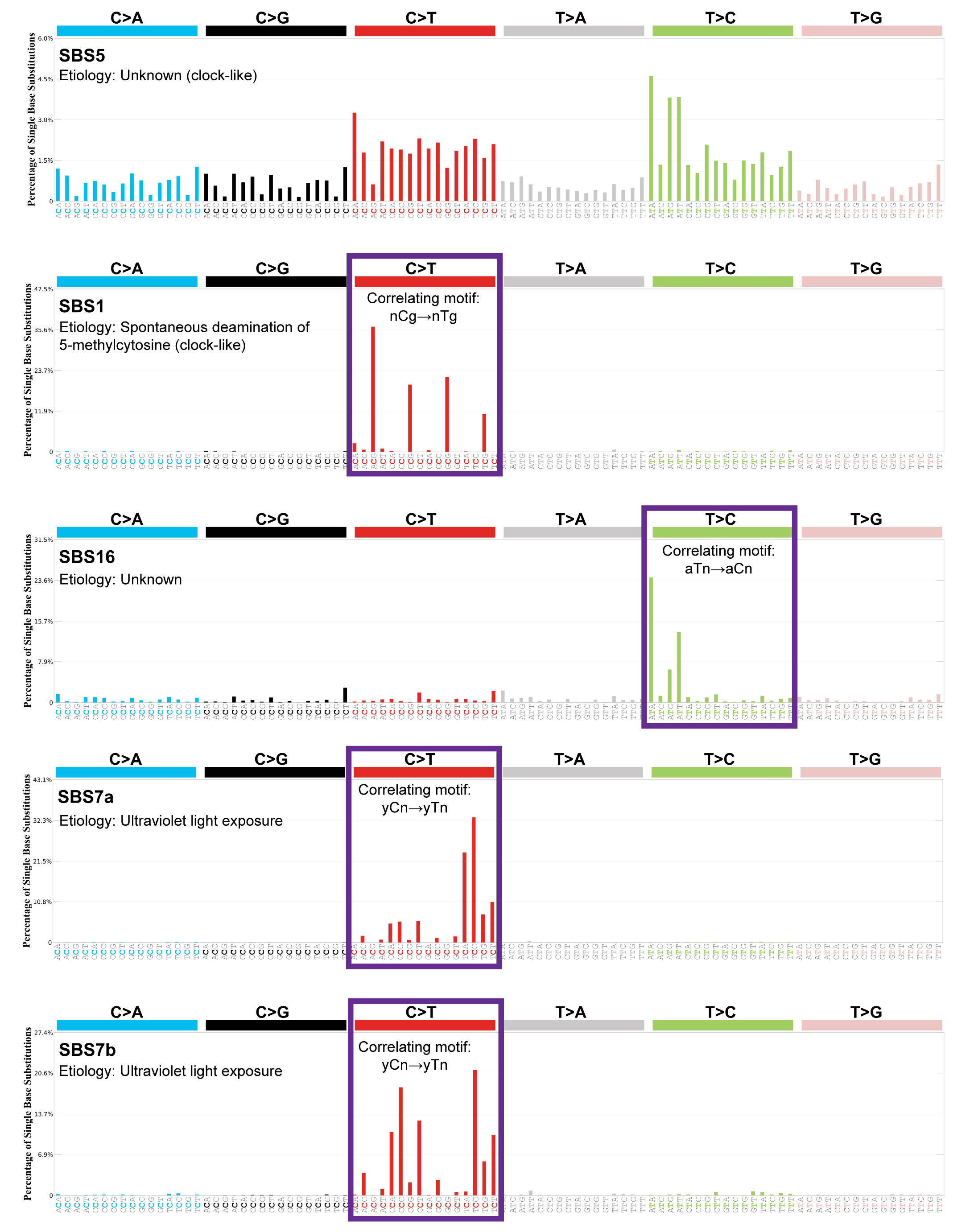

### Fig S5.tif

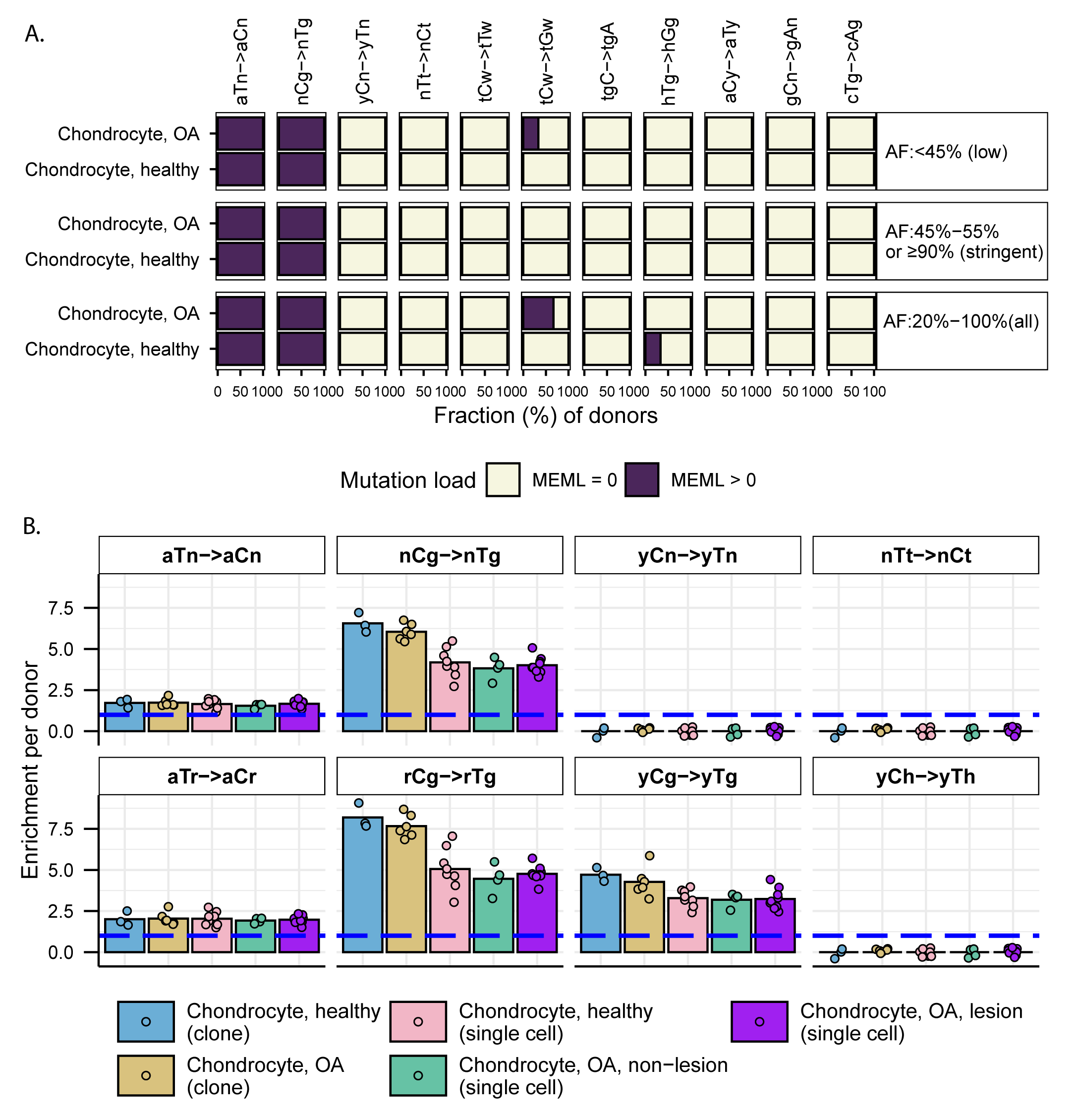

### Fig S6.tif

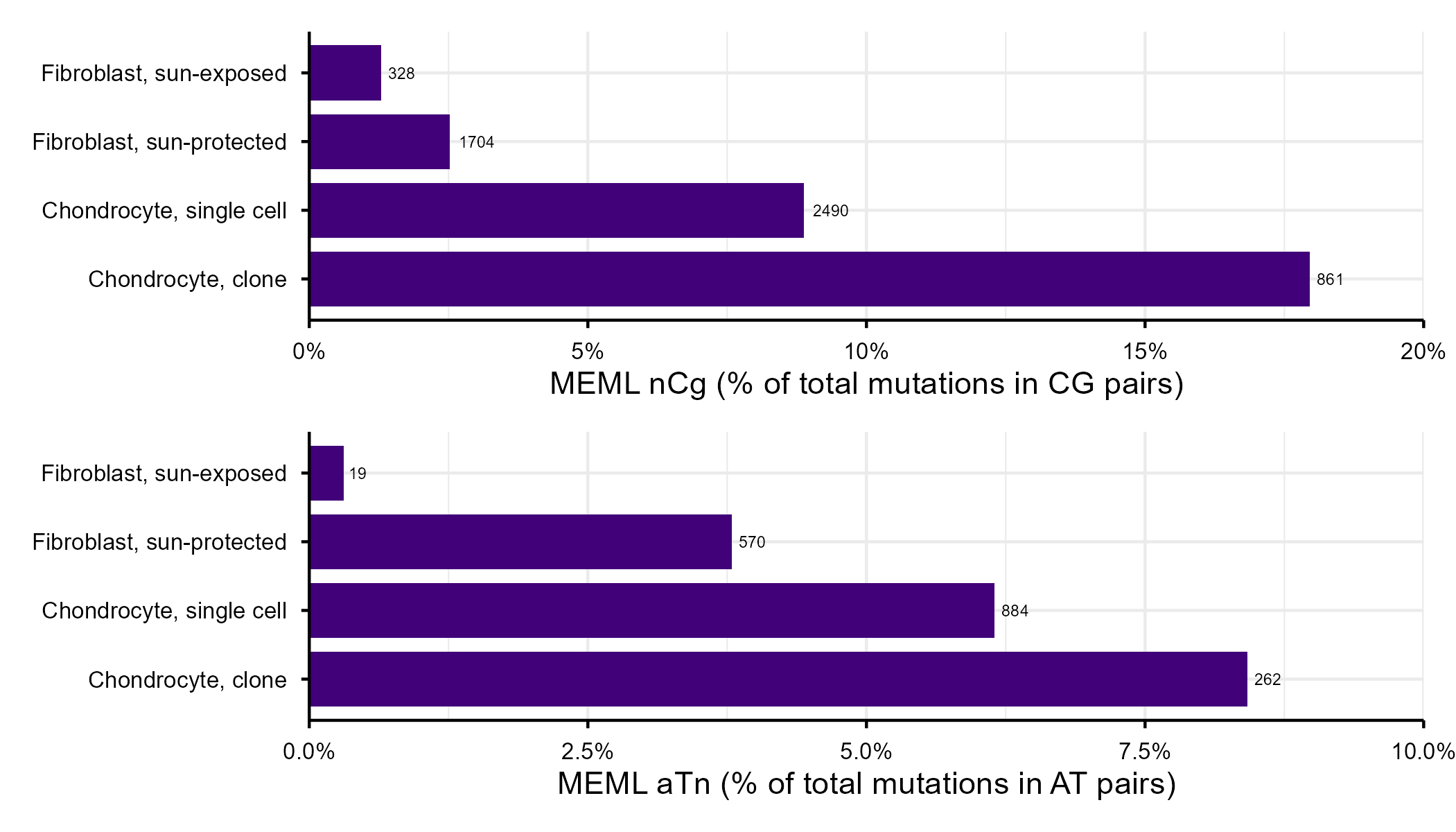

### Fig S7.tif

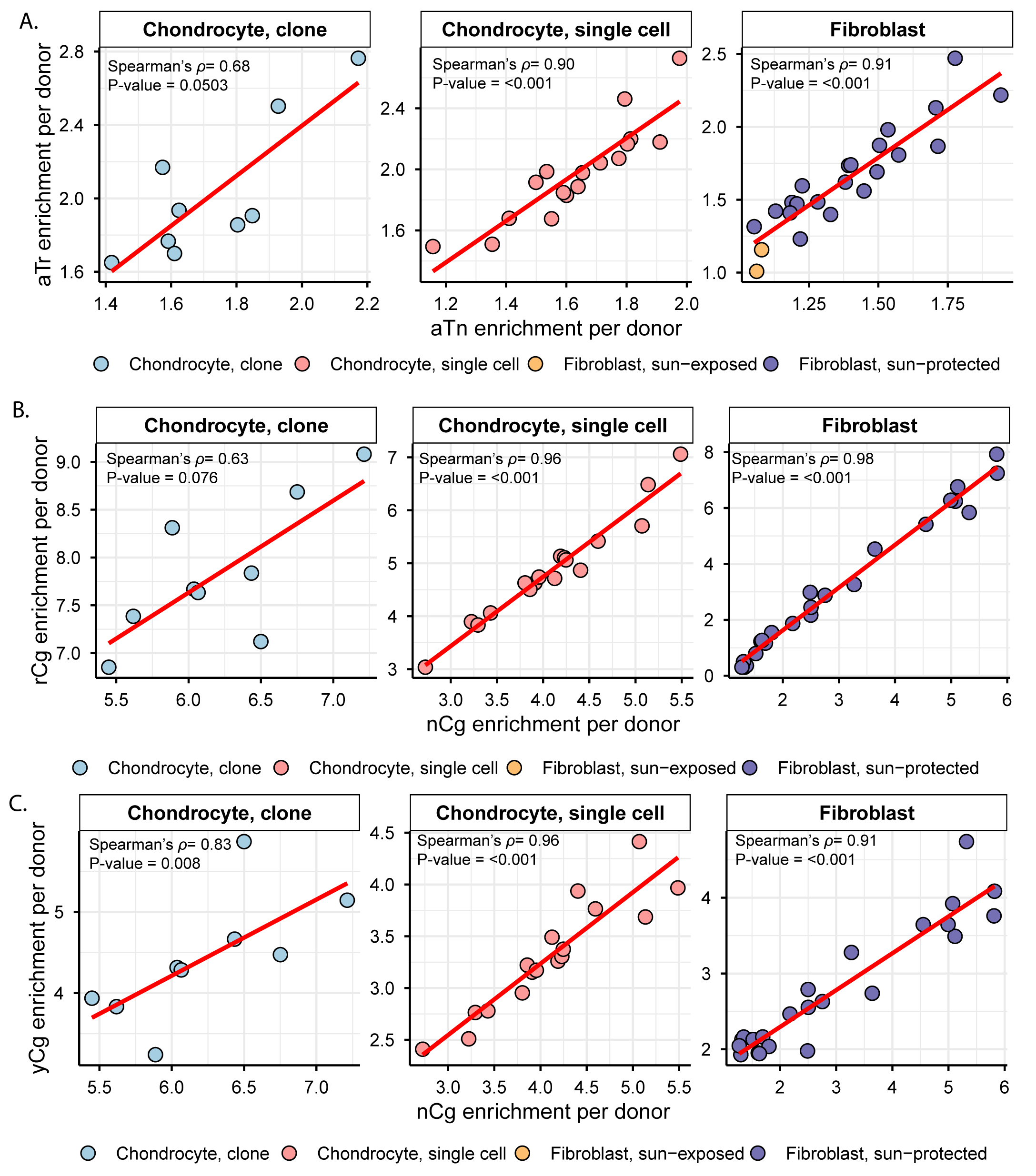

### Fig S8.tif

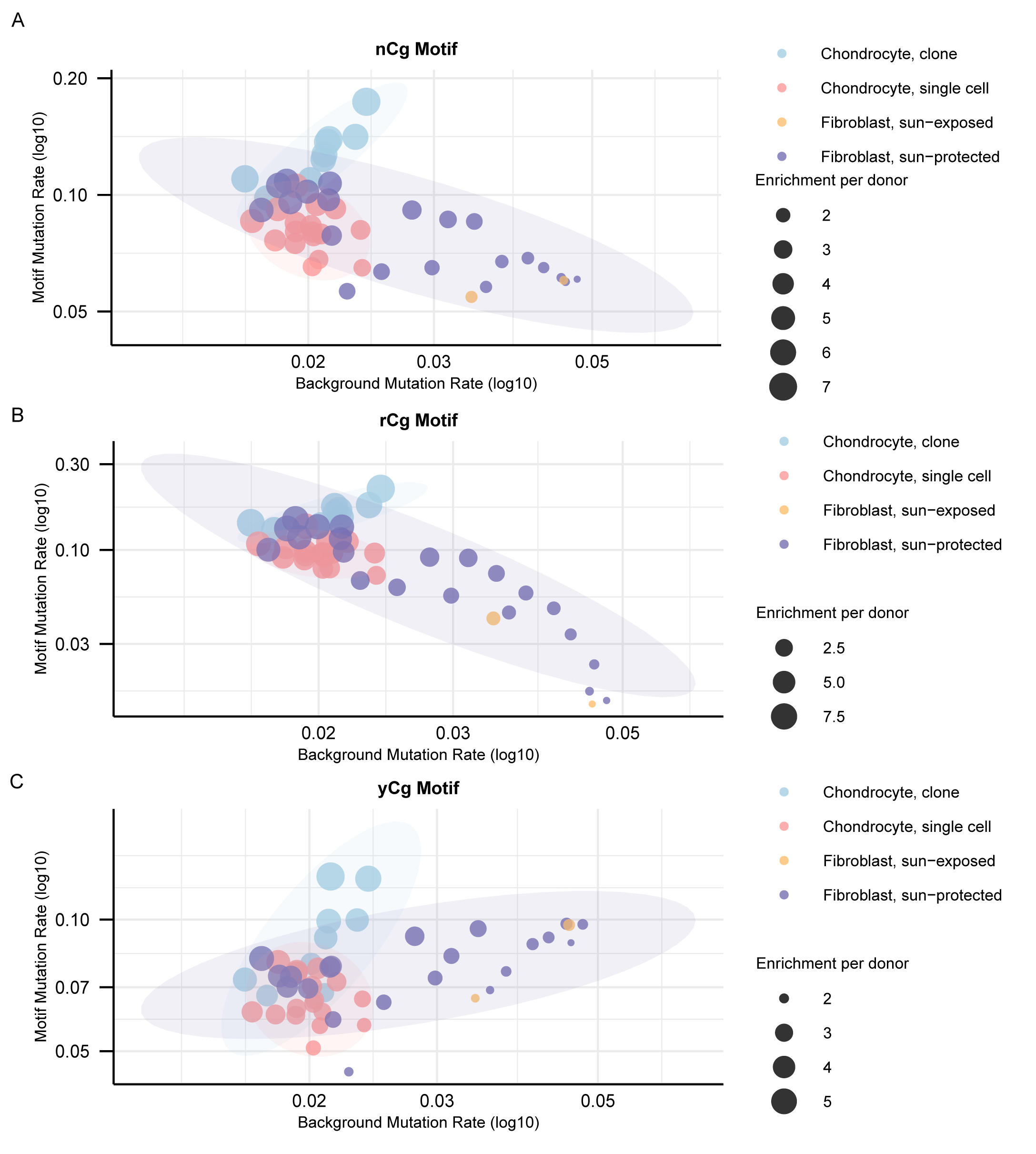

### Fig S9.tif

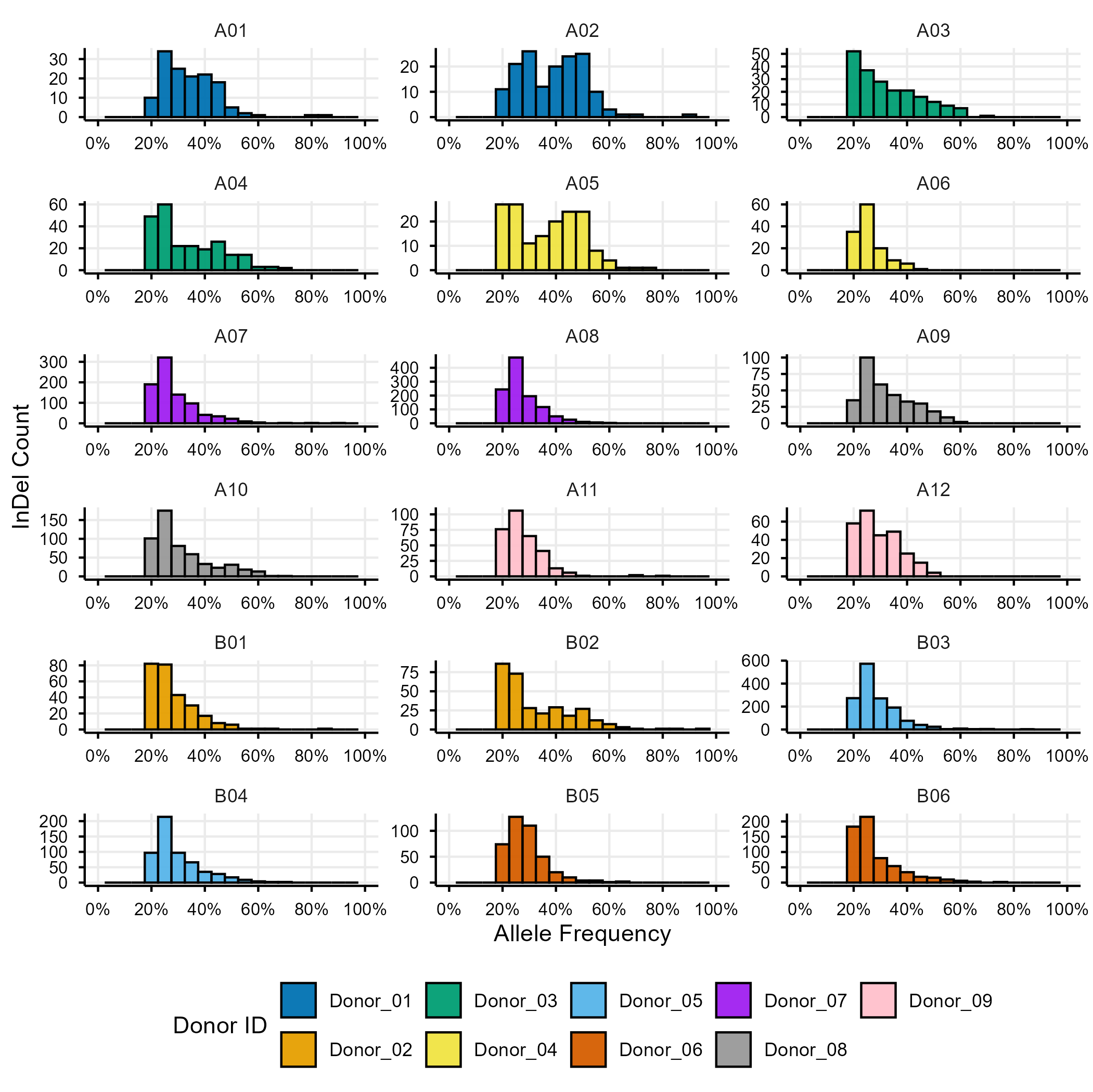

### Fig S10.tif

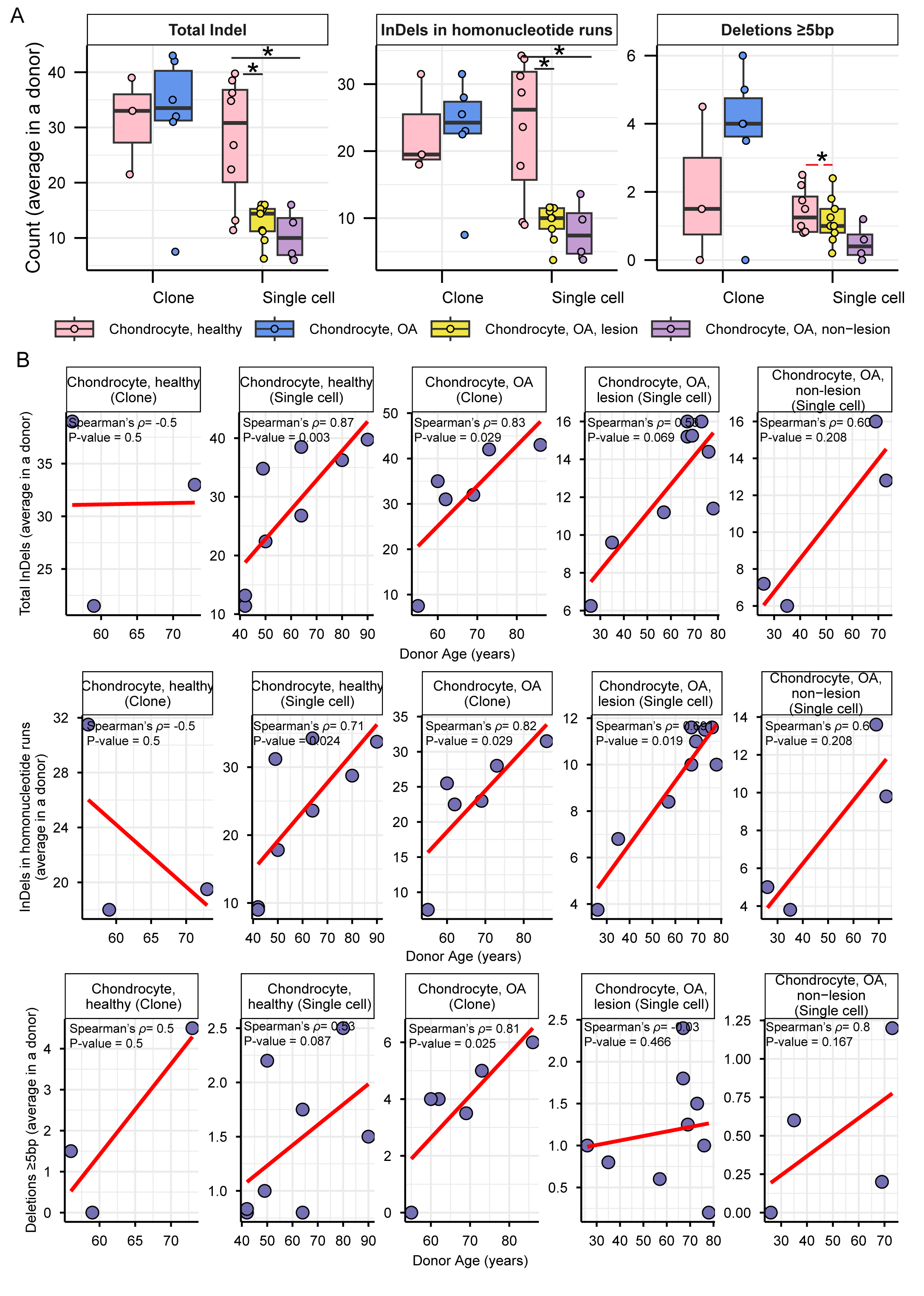

### Fig S11.tif

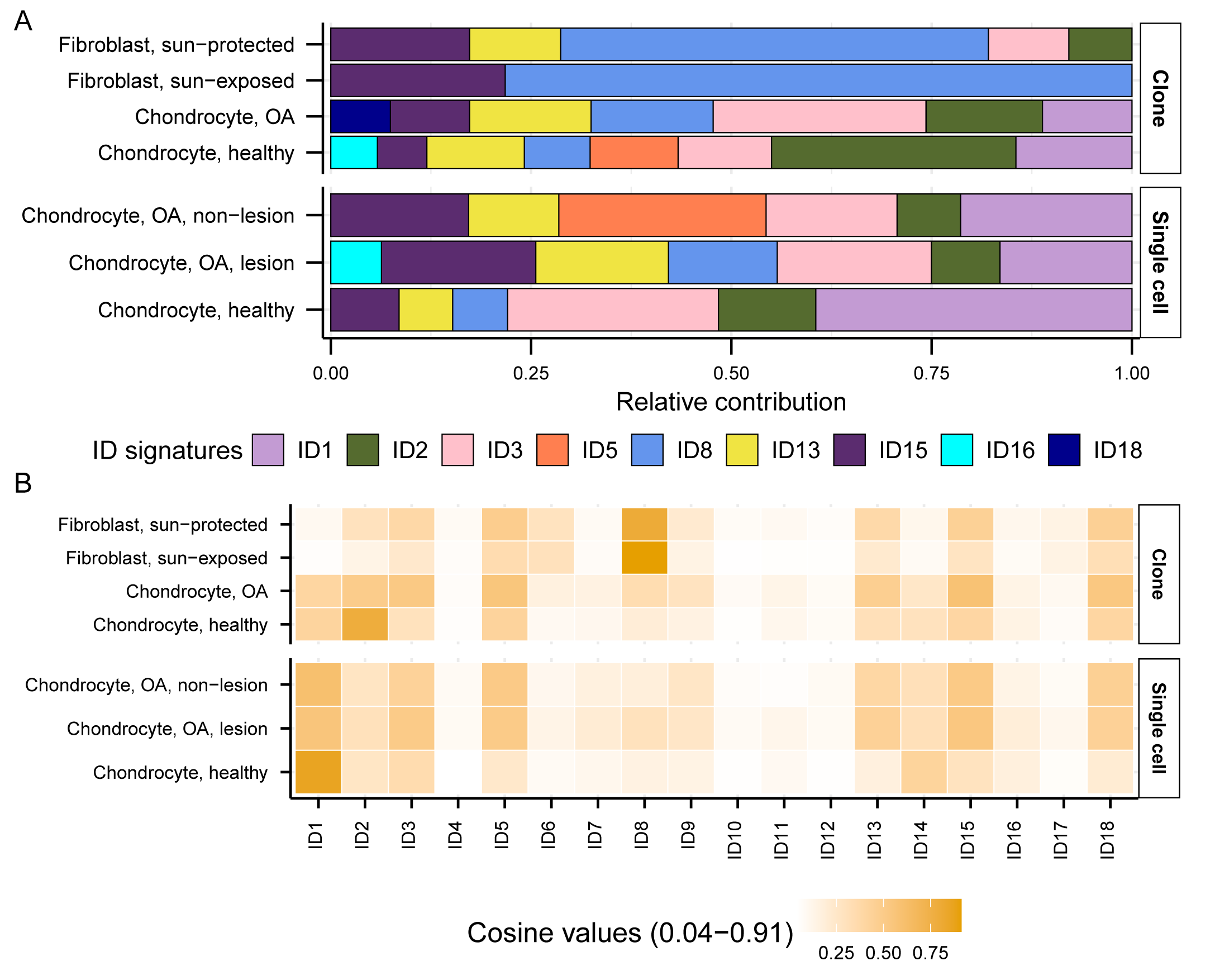
