## Supplemental_tables S1-S7 for "Comparative whole-genome analyses of articular chondrocytes and skin fibroblasts reveal distinct genome instability landscapes in mesenchymal cell types": Supplemental_table_S4.docx

| Reference trinucleotide motif^1^ | Mutant trinucleotide motif^1^ | Motif for correlation with COSMIC SBS signatures | Mutagenic Mechanism | References |
| --- | --- | --- | --- | --- |
| aTn | aCn | aTn🡪aCn | Small epoxide, S_N_2 electrophiles | (Hudson et al., 2023) |
| yCn | yTn | yCn🡪yTn | UV exposure | (Saini et al., 2021; Saini et al., 2016; You et al., 2001) |
| nCg | nTg | nCg🡪nTg | Spontaneous deamination of meCpG | (Pfeifer, 2006; Saini et al., 2021; Saini et al., 2016) |
| nTt | nCt | nTt🡪nCt | UV exposure (minor) | (Saini et al., 2021; Saini et al., 2016) |
| tCw | tTw | tCw🡪tTw | Deamination by APOBEC | (Roberts et al., 2013) |
| tCw | tGw | tCw🡪tGw | Deamination by APOBEC | (Roberts et al., 2013) |
| tgC | tgA | tgC🡪tgA | Redox stress | (Degtyareva et al., 2023)) |
| hTg | hGg | hTg🡪hGg | S_N_2 alkylating agents | (Saini et al., 2020) |
| aTy | aCy | aCy🡪aTy | S_N_1 alkylating agents | (Saini et al., 2020) |
| gCn | gAn | gCn🡪gAn | Acetaldehyde exposure | (Vijayraghavan et al., 2022) |
| cTg | cAg | cTg🡪cAg | Exposure to Aristolactam derived from Aristolochic acid | (Rosenquist & Grollman, 2016) |

Table S4

**List of mutational motifs analyzed in this study.**

1. Letters in motifs indicate IUPAC nucleotide codes.

Degtyareva, N. P., Placentra, V. C., Gabel, S. A., Klimczak, L. J., Gordenin, D. A., Wagner, B. A., Buettner, G. R., Mueller, G. A., Smirnova, T. I., & Doetsch, P. W. (2023). Changes in metabolic landscapes shape divergent but distinct mutational signatures and cytotoxic consequences of redox stress. *Nucleic Acids Res*, *51*(10), 5056-5072. <https://doi.org/10.1093/nar/gkad305>

Hudson, K. M., Klimczak, L. J., Sterling, J. F., Burkholder, A. B., Kazanov, M. D., Saini, N., Mieczkowski, P. A., & Gordenin, D. A. (2023). Glycidamide-induced hypermutation in yeast single-stranded DNA reveals a ubiquitous clock-like mutational motif in humans. *Nucleic Acids Res*, *51*(17), 9075-9100. <https://doi.org/10.1093/nar/gkad611>

Pfeifer, G. P. (2006). Mutagenesis at methylated CpG sequences. *Curr Top Microbiol Immunol*, *301*, 259-281. <https://www.ncbi.nlm.nih.gov/pubmed/16570852>

Roberts, S. A., Lawrence, M. S., Klimczak, L. J., Grimm, S. A., Fargo, D., Stojanov, P., Kiezun, A., Kryukov, G. V., Carter, S. L., Saksena, G., Harris, S., Shah, R. R., Resnick, M. A., Getz, G., & Gordenin, D. A. (2013). An APOBEC cytidine deaminase mutagenesis pattern is widespread in human cancers. *Nat Genet*, *45*(9), 970-976. <https://doi.org/10.1038/ng.2702>

Rosenquist, T. A., & Grollman, A. P. (2016). Mutational signature of aristolochic acid: Clue to the recognition of a global disease. *DNA Repair (Amst)*, *44*, 205-211. <https://doi.org/10.1016/j.dnarep.2016.05.027>

Saini, N., Giacobone, C. K., Klimczak, L. J., Papas, B. N., Burkholder, A. B., Li, J. L., Fargo, D. C., Bai, R., Gerrish, K., Innes, C. L., Schurman, S. H., & Gordenin, D. A. (2021). UV-exposure, endogenous DNA damage, and DNA replication errors shape the spectra of genome changes in human skin. *PLoS Genet*, *17*(1), e1009302. <https://doi.org/10.1371/journal.pgen.1009302>

Saini, N., Roberts, S. A., Klimczak, L. J., Chan, K., Grimm, S. A., Dai, S., Fargo, D. C., Boyer, J. C., Kaufmann, W. K., Taylor, J. A., Lee, E., Cortes-Ciriano, I., Park, P. J., Schurman, S. H., Malc, E. P., Mieczkowski, P. A., & Gordenin, D. A. (2016). The Impact of Environmental and Endogenous Damage on Somatic Mutation Load in Human Skin Fibroblasts. *PLoS Genet*, *12*(10), e1006385. <https://doi.org/10.1371/journal.pgen.1006385>

Saini, N., Sterling, J. F., Sakofsky, C. J., Giacobone, C. K., Klimczak, L. J., Burkholder, A. B., Malc, E. P., Mieczkowski, P. A., & Gordenin, D. A. (2020). Mutation signatures specific to DNA alkylating agents in yeast and cancers. *Nucleic Acids Res*, *48*(7), 3692-3707. <https://doi.org/10.1093/nar/gkaa150>

Vijayraghavan, S., Porcher, L., Mieczkowski, P. A., & Saini, N. (2022). Acetaldehyde makes a distinct mutation signature in single-stranded DNA. *Nucleic Acids Res*. <https://doi.org/10.1093/nar/gkac570>

You, Y.-H., Lee, D.-H., Yoon, J.-H., Nakajima, S., Yasui, A., & Pfeifer, G. P. (2001). Cyclobutane Pyrimidine Dimers Are Responsible for the Vast Majority of Mutations Induced by UVB Irradiation in Mammalian Cells*. *Journal of Biological Chemistry*, *276*(48), 44688-44694. <https://doi.org/https://doi.org/10.1074/jbc.M107696200>
